# Mapfuser: an integrative toolbox for consensus map construction and Marey maps

**DOI:** 10.1101/200311

**Authors:** Dennis van Muijen, Ram K. Basnet, Nathalie N.J. Dek, Chris Maliepaard, Evert W. Gutteling

## Abstract

**Motivation:** Where standalone tools for genetic map visualisation, consensus map construction, and Marey map analysis offer useful analyses for research and breeding, no single tool combines the three. Manual data curation is part of each of these analyses, which is difficult to standardize and consequently error prone.

**Results:** Mapfuser provides a high-level interface for common analyses in breeding programs and quantitative genetics. Combined with interactive visualisations and automated quality control, mapfuser provides a standardized and flexible toolbox and is available as Shiny app for biologists. Reproducible research in the R package is facilitated by storage of raw data, function parameters, and results in an R object. In the shiny app a rmarkdown report is available.

**Availability:** Mapfuser is available as R package at https://github.com/dmuijen/mapfuser under the GPL-3 License and is available for public use as Shiny application at https://plantbreeding.shinyapps.io/mapfuser. Documentation for the R package is available as package vignette. Shiny app documentation is integrated in the application.

## Introduction

Genetic maps play a central role in research and breeding, especially to locate genes or Quantitative Trait Loci (QTL). For several applications it is useful to integrate multiple genetic maps, for example to compare QTLs across populations, to anchor genome assemblies, or to integrate haplotype specific linkage maps in autopolyploids (Bourke et al., 2016). Often in breeding programs it is of interest to integrate genetic information with genomic data. Such a Marey map describing the relationship between centimorgan (cM) genetic distances and physical distances (Chakravarti, 1991) is valuable to estimate local recombination rates and to estimate genetic positions of a given genomic position. The estimation of recombination rates is important for fundamental research to positional cloning of genes, or introgression breeding.

Several software packages are available to construct consensus maps. For example, Joinmap (Van Ooijen, 2006), Mergemap (Wu et al., 2008), and LPmerge (Endelman and Plomion, 2014). To model genomic positions to genetic positions, the MareyMap online can be used (Siberchicot et al., 2017). However, these are all stand-alone tools with different input data formats, consequently requiring researchers spending considerable time on data quality control, back and forth analyses, post-analysis assessment for consensus map construction, Marey map analysis or (final) data visualisation. Reproducible and integrated analyses are ever more important with the barrage of genome assemblies and genetic data becoming available. The mapfuser package and shiny application provide a user-friendly, interactive and modular tool for consensus map construction, interpolation of genetic positions, estimation of recombination rates, and post-analysis assessment. With the implementation of automated quality control features and by integration of different tools, mapfuser opens up the possibility to integrate historical data in present-day research.

## Implementation

The mapfuser R package consists of three major parts: quality control, consensus map construction, and modelling of the relationship between genetic and genomic distance. The map integration depends on the R package LPmerge while for genetic map position and recombination rate estimation we used the R package mgcv to fit thin-plate regression splines (Wood, 2003). In manual data pre-processing errors are difficult to prevent. Therefore, we implemented explicit quality control and data visualisation features. Data can be exported as csv-delimited files or in Joinmap format. A flowchart of the software is presented in Fig. 1.

**Fig. 1.**
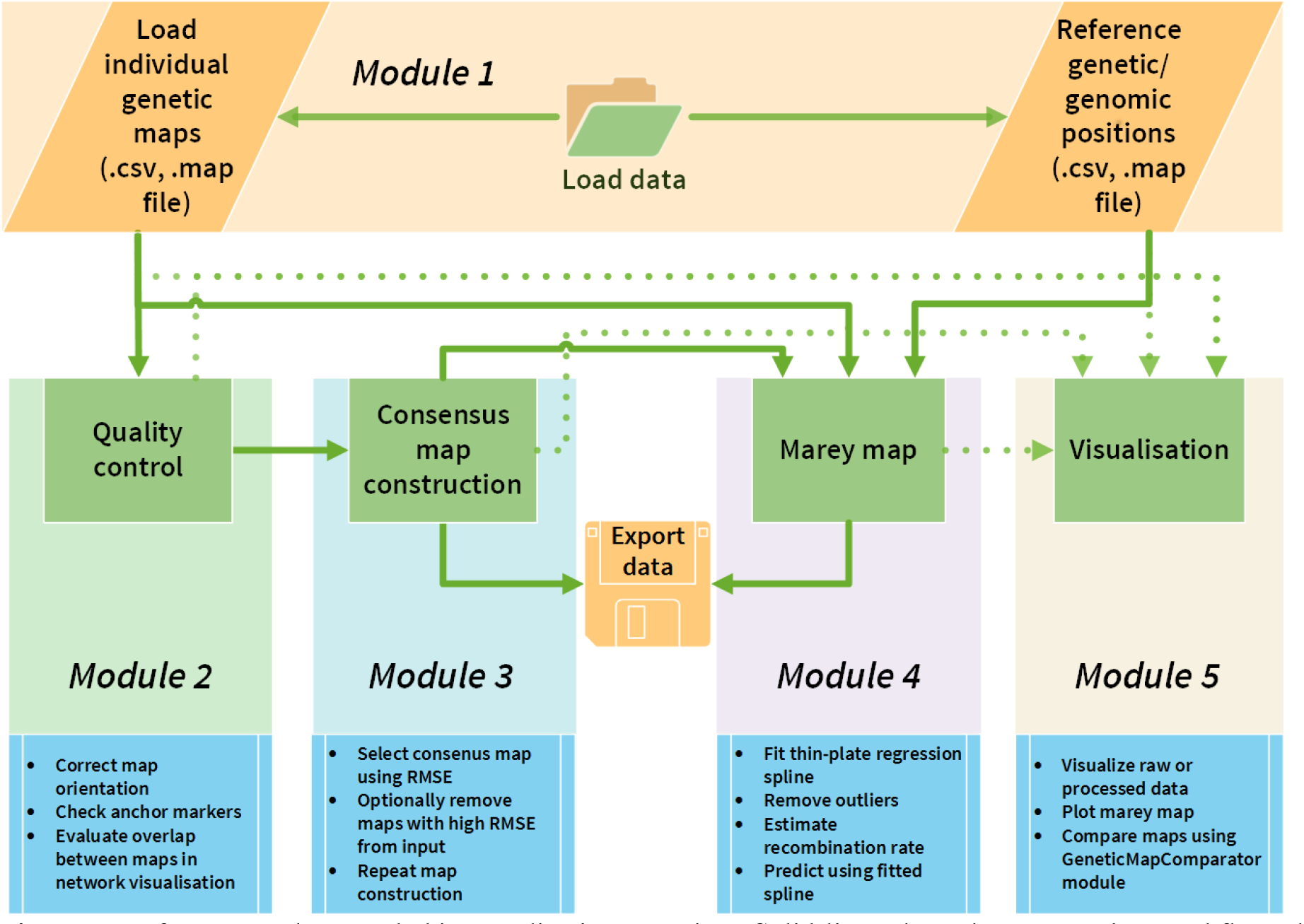
mapfuser R-package and shiny application overview. Solid lines show data processing workflow, dashed lines options for visualisation.

### Module 1 Loading data

Multiple individual linkage maps can be loaded in Joinmap format or in csv-delimited format, with required columns ‘Marker’, ‘Chromosome’, and ‘Position’. For interpolation, a separate csv-delimited file with genome positions is required with columns ‘Marker’, ‘Chromosome’, and ‘Position’ in mega base pairs.

### Module 2 Quality control

The LPmerge package assumes sufficient marker overlap and identical orientation of linkage groups. The map_qc function in the quality control menu filters out linkage groups or entire maps with insufficient anchor markers, as specified by the user. Three markers interspaced at least one cM is considered the minimum requisite to ensure the same orientation of. Input genetic maps are flipped relative to each other first if necessary and subsequently oriented in the direction of a reference map if supplied. Marker overlap between maps is calculated using the igraph R package (Csardi and Nepusz, 2006). The shiny app provides a genetic map visualisation menu adapted from the GeneticMapComparator (Holtz et al., 2017).

### Module 3 Consensus map

With the default setting of “max.interval”- 1:3 as in LPmerge (Endelman and Plomion, 2014), mapfuser gives three versions of integrated maps. The integrated map with the lowest root mean square error (RMSE) is automatically selected, although users can overrule this if needed. In the shiny app, the map integration menu visualizes the integrated map along with summary statistics including the RMSE between individual maps and the consensus integrated map. The consensus map can be exported as csv-delimited file or Joinmap format.

### Module 4 Marey map

To model the relationship between cM and Mbp positions, a minimum of five markers with both a genetic and genomic position is required. Removal of outlier may be warranted due to uncertainties in genetic or genomic maps and is optional in the genphys_fit function. The fitted model can predict the genetic position of unmapped markers or genes based on their genomic positions.

### Module 5 Visualisation

A number of visual diagnostics are available with the generic R S3 plot function: 1) network graphs for each chromosome with edges between maps that provide sufficient overlapping anchor markers 2) a minimum spanning tree based on the network graphs using the number of anchors as edge to build the minimum spanning tree, 3) a scatter plot to compare two maps (input maps, consensus map, reference map, 4) the Marey map depicting the physical distance in mega base pairs vs. the genetic distance in cM along with the fitted thin-plate spline model, 5) a recombination rate plot.

## Case study

Example data provided with the R package contains genetic map data for five Arabidopsis thaliana RIL populations (Simon et al., 2008). Two linkage groups were intentionally flipped and poor quality data was added, which is detected and corrected by the implemented quality control. After construction of a consensus map, a set of PCR markers from a sixth A. thaliana population (Loudet et al., 2002) was interpolated towards the consensus map. Ordinary least squares regression indicates that the estimated positions fit well for each chrome (R2 > 0.99) relative to the true genetic map positions and compares well with estimates from MareyMap Online. The subtle difference between both methods is that the penalized spline is more robust for local deviations. Single positions on a genetic map are affected by Mendelian sampling errors, genotyping errors, and uncertainties in statistical inference in ordering and distance estimation. However, the result is that possible biologically significant differences in recombination rate, represented by one or very few data points, are neglected. Nonetheless, the fit in terms of R-squared is better using a penalized approach in the example data.

## Acknowledgements

We are grateful to Bart Lamboo for shiny server troubleshooting and thank Marcel Kerkveld for consultation on software architecture.

